# C1orf109L promote R-loop accumulation induced DNA damage to inhibit cell growth

**DOI:** 10.1101/625749

**Authors:** Dou Peng, Li Yiqun, Xie Wanqiu, Zhang Xiaoqing, Zhang Dandan, Ci Yanpeng, Zhang Xiaohan, Qiao Shupei, Muhammad Luqman Akhtar, Han Fang, Yu Li

**Affiliations:** School of Life Science and Technology, Harbin Institute of Technology, Harbin City, China

**Keywords:** C1orf109L, R-loop, cell cycle arrest, DNA damage, ubiquitination

## Abstract

As a function unknown gene, C1orf109 is lower expression in various cells. Here, we reported that C1orf109L, the longest variant of C1orf109, which interacted with R-loop-regulating proteins to trigger R-loop, a three-stranded nucleic acid structure frequently mediated genome instability, accumulation. C1orf109L induce chronic DNA damage to promote P21 upregulation and strongly inhibits cell growth *in vitro* and *in vivo* by arresting the cell cycle in the G2 phase. With camptothecin (CPT), an R-loop activator, treatment, C1orf109L further triggers R-loop accumulation-induced DNA damage and promotes cell death by activating cell-death pathway. Furthermore, CPT treatment increases C1orf109L ubiquitination and turnover, which inhibits cell death and promotes the G0/G1 phase of the cell cycle. Therefore, our data illustrated the mechanisms underlying C1orf109L-related cell growth inhibition and provide feasibility and limitations for C1orf109L as a potential target for cancer therapy.

---

Cell proliferation depends on the cell cycle. In abnormal cells, such as benign tumor cells and cancer cells, cell-cycle dis-regulation promotes cell growth [1]. For the cancer therapy strategies are mainly to kill cancer cell or inhibit cancer cell growth. Inducing cancer cell DNA damage is an efficient way to induce cancer cell death or growth inhibition. When genomic DNA is damaged, many pathways involved in cell-cycle regulation are activated, the most important of which involves activation of the DNA-damage checkpoint to arrest the cell cycle at different stages, including G1, S, and G2/M, which ensures that the damaged cell has enough time to repair DNA damage [2,3]. If DNA damage is so severe or the rate of damage is faster than the rate of the repairing process, then the cellular-death pathway is activated [4].

Among the DNA-damage checkpoints, the G1 and G2/M checkpoint restricts mitosis onset in response to multiple exogenous and endogenous factors [5]. With respect to endogenous factors, several proteins act as RNA-processing regulators that control the cell’s fate, while dysfunction of RNA-processing-related proteins could trigger R-loop accumulation, and RNA–DNA hybrids can lead to DNA damage and induce-cell cycle arrest, which is an important factor for genome instability in abnormal cells, especially in cancer cells [6,7]. The R-loop also controls gene expression [8] and immunoglobulin (Ig)G class switching [9]. However, the regulation of R-loop proteins is poorly explored. Thus, in this study, we identified the novel gene, *c1orf109*, which is downregulated in a variety of cell lines. C1orf109L-eGFP expression triggers R-loop accumulation to induce DNA damage and arrest the cell cycle.

We report that C1orf109L interacted with R-loop associated proteins and located on R-loops. C1orf109L overexpression that triggered R-loop accumulation no matter with or without CPT treatment in cells. The accumulated R-loops induced DNA damage which inhibits cell growth in vitro and in vivo. In terms of the mechanism underlying cell growth, the C1orf109L expression activated the cell-cycle checkpoint controlling the mitotic-control protein P21 (encoded by *cdkn1a*) and inactivated phosphorylation CDK1 accumulation which lead to an aberrant G1/S and G2/M transitions in cells. This process does not involve the R-loop or the C1orf109L gene-regulation function, as based on analysis of global gene-expression profiling. C1orf109L triggered further R-loop accumulation and promoted cell death in response to CPT treatment. C1orf109L induced DNA damage was enhanced and cell death pathway was activated by CPT treatment. However, with CPT treatment, C1orf109L has been ubiquitinated and degraded. Meanwhile, the cell cycle enters into the G0/G1 phase to inhibit CPT-induced cell death. In summary, our study uncovers C1orf109L as an R-loop regulator to promote DNA damage and activate DNA-damage checkpoints. We also revealed the cell-feedback mechanism that protects cells from C1orf109L-eGFP induced excessive DNA damage in response to CPT.

## Result

### The C1orf109L protein–protein interaction network

To explore the function of C1orf109L in cells, we need to know the interaction proteins of C1orf109L, therefore we sought to identify C1orf109L-interacting proteins by tandem mass spectrometry-based affinity proteomics, using Flag-tagged C1orf109L immunoprecipitated from HeLa cells. There are 236 specific interaction proteins (Fig 1A) and functions that are mainly rich in RNA metabolism and processing, as determined through Gene Ontology (GO) annotations (Fig 1B). To further verify that C1orf109L-eGFP can bind with RNA-related proteins, we found that C1orf109L-eGFP in the chromosome was reduced by different concentrations of RNAse A, which digested the chromosome of HEK-293T cells (Fig 1C). Network clustering identified highly interconnected nodes within the network, possibly reflecting protein complexes that may interact with C1orf109L-eGFP within the context of distinct cellular processes. Enrichment analysis based on GO annotations indeed showed that these sets of C1orf109L-interacting proteins function in RNA processing, RNA metabolism, and genome structure maintenance (Fig 1D and 1E). We selected interactors from 31 interactors and, as confirmed through immunoprecipitation, we found that C1orf109L-eGFP can bind with DHX9 and NMP1 (Fig 1F).

### C1orf109L binding with R-loops and promote DNA damage

According to László Halász et al., who identified R-loop-related proteins in HeLa cells [10] and our results showed that C1orf109L-eGFP can bind with many R-loop-regulating proteins to promote cell-format R-loops, such as DHX9, an RNA helicase that is closely related to R-loop formation. Loss of DHX9 can promote R-loop formation and enhance R-loop-induced DNA damage in response to CPT, a topase I inhibitor, which can promote R-loop formation by stopping TOP I and promoting Pol II striking with TOP I [11]. We found that among 31 interactors of C1orf109L, 16 interactors were R-loop-related proteins (Fig 2A); moreover, we confirmed that C1orf109L-eGFP can bind with the R-loop in an RNA-dependent manner, we isolated the HeLa cell nucleus and extracted R-loops by immunoprecipitation with or without RNAse A treatment. The results showed that C1orf109L-eGFP reduced upon RNAse A treatment, as compared with no RNAse A treatment (Fig 2B and C).

R-loop regulation proteins abnormal will cause R-loop accumulation, so we measured the number of cells displaying fluorescence intensity of the nucleus via R-loop detection using an R-loop specific antibody (S9.6) during immunofluorescence. The results demonstrated that C1orf109L-eGFP expression can significantly (*p*<0.001) increase the fluorescence intensity of the R-loop in the nuclei (Fig 2D). However, when RNAseH1-eGFP, an R-loop digestion enzyme, is overexpressed, the fluorescence intensity of the nuclei had significantly (*p*<0.001) decreased (Fig 2D). CPT is an anticancer drug which is also a R-loop activator. During the immunofluorescence assay, we observed that C1orf109L-eGFP expression can significantly enhance the fluorescence intensity of the nucleus R-loop in response to CPT (Fig 2E). we also discovered that C1orf109L-eGFP (green) present colocalization (black arrows) with the R-loop (red), furthermore, in nuclear, large-scale colocalization for C1orf109L and R-loops in response to CPT (Fig 3A).

R-loop accumulation is an important reason for DNA damage, During karyotype analysis, C1orf109L-eGFP promoted the cell to form an uncompressed chromosome that belongs to the G2 phase of the cell cycle. Interestingly, we found a breakage point on the uncompressed chromosome (indicated by the arrow; Fig 3B). To verify whether C1orf109L-eGFP can trigger DNA damage, a comet assay was conducted, and the results showed that the tail of the DOX+ group was remarkably longer than that of the DOX- group (Fig 3C). Moreover, with CPT treatment, the expression of C1orf109L caused heavy DNA damage. The results illustrated that C1orf109L-eGFP expression can cause DNA damage (Fig 3C). CPT induced R-loop accumulation, which promoted γH2AX sustained upregulation for 240 minutes; following this, there was a rapid drop [12,13]. Based on our findings, it was evident that C1orf109L-eGFP participated in RNA processing and triggered R-loop accumulation and caused serous DNA damage in response to CPT. We supposed that C1orf109L-eGFP can promote DNA damage and sustain γH2AX over a long period of time. we performed HeLa-Tet-on cells were treated with or without DOX for 24 hours and treated with CPT for the indicated amounts of time; the results showed that γH2AX increased gradually over the course of CPT treatment when compared to the DOX- group, which demonstrated that γH2AX increased gradually until 4 hours of CPT treatment, at which point it dropped rapidly until 6 hours (Fig 3D). The results indicated that C1orf109L-eGFP expression can cause severe DNA damage in response to CPT.

### C1orf109L-eGFP expression inhibits cell growth in vitro and in vivo

DNA damage is an efficient way to inhibit cell growth, for above result, C1orf109L may be a powerful target for some disease therapies, such as cancer. In order to identify the expression of C1orf109 in various cells, we generated a monoclonal antibody that can specifically recognize different tags of exogenous C1orf109L and C1orf109S via immunoblotting (Fig EV1A and B). After verifying C1orf109 antibody, we performed immunoblotting to identify the C1orf109 protein level in many types of cell lines. However, we cannot detect C1orf109 protein expression in indicated cells (Fig EV1C). A previous study showed that C1orf109 expression was not observed in various types of abnormal cells, such as keloids, mainly because of hyper-methylation in the promoter region of the C1orf109 gene[14]. Thus, we hypothesis that undetectable C1orf109 expression is caused by epigenetic involvement. To verify whether epigenetic modification of C1orf109 regulates C1orf109 expression, the C1orf109 protein and mRNA levels were detected following treatment with different concentrations of DNA methylation enzyme inhibitor, 5’-Aza-deoxycytidine (5-aza), or histone deacetylation enzyme inhibitor, Trichostatin A (TSA), in HeLa cells, as indicated over time. Strikingly, TSA treatment can improve C1orf109 expression, as indicated by the observed concentration at 24 hours (Fig EV1D). Thus, we detected the mRNA and protein levels of C1orf109 following TSA treatment at different times with working concentrations. We found that C1orf109 expression was markedly improved in a time-dependent manner under TSA treatment (Fig EV1E and F).

To explore whether C1orf109L can inhibit cancer cell growth as a potential therapy target, we perform the transfection to over-express C1orf109L in various cancer cell lines and immortalized cells, the result show that C1orf109L expression can significant inhibit cell growth (Fig EV1G). In order to futher to explore the function of C1orf109L, we established a reliable doxycycline (DOX)-inducible common model using the HeLa-Tet-on and HEK-293-Tet-on cell lines, which expressed eGFP-tagged C1orf109L induced by DOX (Fig EV1H). To explore the influence of HeLa under the expression of C1orf109L-eGFP, we treated HeLa cells with DOX (DOX+) or dimethyl sulfoxide (DMSO; DOX-), as indicated over time and as observed in the HeLa cell phenotype over an 8-hour time lapse. We found that C1orf109L-eGFP expression (DOX+ group) can reduce the numbers HeLa cells undergoing cell division when compared with the number of cells in the DOX- group (Fig 4A; red arrows indicate cell division and movie1 is DOX- group and movie2 is DOX+ group). These results indicate that cell growth was inhibited under C1orf109L-eGFP expression. To further confirm whether C1orf109L-eGFP expression inhibited cell growth, we analyzed the clone-formation ability of cells under the expression of C1orf109L-eGFP, which showed that the expression of C1orf109L-eGFP can significantly (*p*<0.001) reduce clone numbers in both HeLa and HEK-293 cells (Fig 4B), as compared with the induction of eGFP expression (Fig EV2A). Meanwhile, we analyzed the cell growth rate in the DOX+ and DOX- groups using Cell Counting Kit-8 (CCK-8). The results illustrated that C1orf109L-eGFP can strongly (*p*<0.001) inhibit cell growth both in HEK-293 and HeLa cells (Fig 4C) when compared with the induction of eGFP expression (Fig EV2B). R-loops causing DNA damage can activate the DNA damage signaling pathway to inhibit cell growth [15,16]. It is important to assess whether C1orf109L-eGFP can cause R-loop formation, as it may represent a major reason for cell growth inhibition; therefore, to analyze this, the clone-formation assay demonstrated the same phenotype, highlighting how RNAseH1-eGFP overexpression can reverse C1orf109L-eGFP expression, causing clone-formation inhibition (Fig 4D). Further, we overexpressed RNAseH1-eGFP in the DOX+ and DOX- groups and measured the cell-growth rate using CCK-8. The results indicated that RNAseH1-eGFP was overexpressed in C1orf109L-eGFP-expressed cells, which can partially reduce C1orf109L-eGFP, causing cell-growth inhibition (Fig 4E). These results indicated that C1orf109L-eGFP can inhibit cell growth by promoting R-loop accumulation.

Based on the aforementioned results, C1orf109L-eGFP exhibited strong cell-growth inhibition capabilities in vitro. To investigate whether C1orf109L-eGFP has potential ability in anti-tumor growth, we performed a nude-mouse tumor-transplantation experiment to assay tumor growth inhibition under C1orf109L-eGFP expression. We inoculated DOX-inducible HeLa cells subcutaneously into the lower flank of 6-week-old nude mice. Two weeks later, following tumor formation, the mice were divided into one of two groups according to similar tumor size. Mice were fed with or without DOX containing water and the tumor volume was measured using a digital caliper every 2 days (Fig 4F). After 12 days, the mice were sacrificed. The results demonstrated that the tumor volume of mice in the DOX+ group was smaller than that in the DOX- group (Fig 4F and 4G). The tumor growth rate also presented the same trend (Fig 4H) and the tumor weight in the DOX+ group was remarkably lighter (*p*<0.01) than that in the DOX- group (Fig 4I); However, the stable expression of the eGFP tumor (DOX+ group) highlighted how tumor volume, the tumor growth rate, and the tumor weight did not demonstrate significant differences when compared with those of the DOX- group (Fig EV2E and 2F). These results show that C1orf109L-eGFP strongly inhibited cell growth both in vitro and in vivo.

### C1orf109L-eGFP influences the cell-cycle G1/S phase transformation and arrests the cell cycle in G2 phase

Cell proliferation inhibition is caused by cell cycle arrest and DNA damage is an efficient and rapid way to arrest the cell cycle by activating the cell-cycle checkpoint [17,18]. To test cell cycle arrest which cell cycle phase, we performed HeLa-Tet-on cells were treated with TDR for 16 hours, and TDR was then withdrawn for 10 hours. Combining treated with TDR, and with or without DOX, was performed to synchronize the cell cycle at the G1/S boundary (Fig EV3A). Intriguingly, when TDR treatment was withdrawn, the cells were not fully arrested; rather, they progressed very slowly into S phase in the DOX+ group, as compared to the DOX- group (Fig EV3A). The R-loop can also arrest the cell cycle in the G2/M phase by activating the cell-cycle checkpoint [19]. For that reason, we performed flow cytometry to detect the cell cycle to analyze the effects of the cell cycle under C1orf109L-eGFP expression. Compared to the control group (DOX-), the HeLa and HEK-293 cells in the DOX+ group exhibited a significant increase in the number of cells in the G2/M phase (*p*<0.001; Fig 5A). For HeLa cells, the cell cycle time is clear, for cell cycle at M phase, cell shape become spherify, which is the difference between that G2 phase HeLa cells(Fig 5B). To further identify whether C1orf109L-eGFP arrested the cell cycle in G2 or M, the cells were treated with nocodazole, an M-phase inhibitor; meanwhile, the cells were treated with DOX. The results demonstrated that in the control group cells (DOX-/nocodazole+), about 95% of cells entered the M phase with chromosome condensation; however, the number of cells in the M phase in the C1orf109L-eGFP-expressed group (DOX+/nocodazole+) were significantly reduced and about 30% of cells entered into the M phase, as compared to the control group. Therefore, nocodazole treatment can reduce the number of cells in the M phase with C1orf109L-eGFP expression (Fig EV3B). The same phenotype was observed in HEK-293 cells (Fig EV3C). These results provided evidence in support of the idea that cells in the DOX+/nocodazole+ group feature tetraploid DNA without the M-phase phenotype, which indicates that C1orf109L-eGFP arrests the cell cycle in the G2 phase.

C1orf109L-eGFP expression in a short time so that it did not have enough time to regulate the expression of other genes. Therefore, we arrested the cell cycle at the G1/S boundary, as described earlier, and cell cycle released with or without C1orf109L-eGFP expression in about 12 hours. Interestingly, the cell cycle appeared to be blocked from the G2/M (4N) to G1 (2N) phase, releasing at 10 hours (Fig 5C). We performed another experiment to arrest the cell cycle at the G1/S boundary using the same treatment as above, and released it with or without C1orf109L-eGFP expression in about 12 hours under nocodazole treatment. We found that C1orf109L-eGFP expression can remarkably reduce 50% of M-phase cells when compared to the control group in nearly 100% of M-phase cells (Fig 5D). We exclude R-loop caused gene expression change by analyzed the transcriptome of inducing C1orf109L-eGFPfor 12 hours and 24 hours in HeLa cells. the result showed that the expression of C1orf109L do not change cell cycle related gene expression through GO annotion (Fig EV4A, B and C). Furthermore we conformed C1orf109L do not change G2-M transtration gene change by Releasing the cell cycle and C1orf109L-eGFP expression about 8 hours, as described, we detected the RNA and protein levels of those genes that directly control the cell cycle for the G2/M transition, and we found that CDK1 and CCNB1 expression did not alter either the RNA or protein levels (Fig EV4D and E). These results demonstrated that C1orf109L-eGFP arrested the cell cycle in the G2 phase.

We detected DNA damage marker γH2AX and found that the level of γH2AX in the DOX+ group was elevated when compared with that of the DOX- group, both in the HeLa and HEK-293 cells (Fig 5E). Damaged DNA can lead to the downstream activation of the cell-cycle checkpoint; cell-cycle checkpoint protein P21 belongs to the downstream DNA damage pathway and is a cell-growth inhibitor that controls the cell cycle G1-S and G2-M phases. Moreover, its upregulation can also cause cell senescence [20,21]. We found that C1orf109L-eGFP expression can significantly upregulate P21 protein levels and showcased how P21, found downstream of the CDK1-inactivated phosphorylation forms, was significantly upregulated (Fig 5E). To test whether C1orf109L-eGFP caused P21 upregulation and thus led to cell-growth inhibition, we knocked down P21 under C1orf109L-eGFP expression (Fig 5F); we found that p21 downregulation can significantly reverse cell-growth inhibition (Fig 5F). The results showed that C1orf109L-eGFP caused DNA damage and activated the cell-cycle checkpoint.

### C1orf109L-eGFP promotes R-loop-dependent DNA damage and cell death in response to CPT

In above result CPT will caused serious DNA damage with expression of C1orf109L and promote γH2AX upregulation graduation with time dependent. We performed a time lapse to observe the phenotype of the expression of C1orf109L-eGFP in response to CPT; as such, we seeded the same number of DOX-induced RFP expression HeLa as control cells (red cells) and DOX-induced C1orf109L-eGFP HeLa cells (green cells) on a plate and induced expression for 24 hours; time lapse was detected over 8 hours (Fig 5A). We observed that the green cells (C1orf109L-eGFP expression) began to die after about 5 hours of CPT treatment, and the red cells (RFP expression) did not die until 8 hours (Fig 6A and movie3). HeLa-Tet-on cells expressed C1orf109L-eGFP in response to CPT within 12 hours; when we measured the cell number, the results showed that the number of cells in the CPT+/DOX+ group had reduced about 50% when compared with the cells in the CPT-/DOX+ group. Moreover, there was no difference between the number of cells in the CPT-/DOX+ group and CPT-/DOX- group (Fig 6B). The results indicated that C1orf109L-eGFP enhanced cell death in response to CPT. C1orf109L-eGFP expression can cause cell death with CPT treatment, so we preformed Western blot analysis to analyze the protein levels along the cell-death pathway. During this analysis, HeLa-Tet-on cells were treated with or without DOX for 24 hours and then treated with CPT for the indicated amount of time; the results showed that under CPT treatment, C1orf109L-expression could active caspase-9 and PARP1 was cleaved (Fig 6C). These results indicated that C1orf109L could trigger R-loop accumulation, induce DNA damage, and activate the cell-death pathway in response to CPT.

### C1orf109L-eGFP degrades cells through an ubiquitinated proteasome pathway in response to CPT

Based on the aforementioned discovery that C1orf109L-eGFP exhibits weakened fluorescence in response to CPT, we assumed that there was a mechanism underlying the action of CPT to promote the degradation of C1orf109L-eGFP to ensure cell survival. Therefore, we assessed the stability of C1orf109L-eGFP with CPT treatment. The results showed that the protein level of C1orf109L gradually decreased with prolonged treatment times with CPT. Further, we found that the additional treatment of MG132, a proteasome inhibitor, on cells could hinder the decrease in protein levels of C1orf109L-eGFP (Fig 7A). These results suggest that C1orf109L is degraded by the proteasome pathway in response to CPT. Previous studies have found that C1orf109L can be degraded by the ubiquitinated proteasome pathway (Fig EV5A, B and C), so we hypothesized that C1orf109L ubiquitination can be promoted by CPT. The results also showed that the C1orf109L ubiquitination level in the CPT-treated group was significantly higher than that in the DMSO group (Fig 7B). Furthermore, we analyzed the ubiquitination linkage of C1orf109L by constructing different ubiquitin molecular mutants. The results indicated that K6-, K11-, and K33-ubiquitin linkages of C1orf109L-eGFP were upregulated in response to CPT (Fig. 7C).

When the cell suffers from unfavorable factors, such as a lack of nutrition, cytotoxicity, and DNA damage, the cell will enter into a dormant phase (G0/G1 phase) to protect itself from unfavorable factors [22,23]. We analyzed whether the cells enter dormancy (G1/G0 phase) after complete degradation of C1orf109 under long-term CPT treatment. We found that the long-term treatment of cells with CPT, particularly following the expression of C1orf109L-eGFP for 24 hours (CPT+/DOX+), resulted in more G0/G1 phase cells than did those cells that underwent a lengthy treatment of CPT but did not express C1orf109L-eGFP (CPT+/DOX-; Fig. 7D). These results suggest that C1orf109L-eGFP will be degraded by the ubiquitinated proteasome pathway under CPT treatment, after which point the cells will enter into the G0/G1 phase. These data, together with the results of our ubiquitination experiment, establish the fact that in the presence of CPT, C1orf109L-eGFP had degraded while the cells entered into G0/G1 phase to avoid cell death.

*C1orf109* is an uncharacterized gene, although the shortest variants of *C1orf109* were reportedly involved in cell growth [24]. Previous research has failed to consider the mechanisms underlying cell-cycle regulation, and there are no reports on the function of C1orf109L, the longest variant of *C1orf109*. Our work showed that C1orf109L-eGFP strongly inhibited cell growth *in vitro* and *in vivo*. In terms of the mechanism underlying cell-growth inhibition, we detected C1orf109L-eGFP-interaction proteins that are primarily involved in RNA processing. The RNA processing of protein dysfunction will cause cell-growth inhibition via regulation of the transcription process or genome instability including telomere instability [25,26]. One of the most notable growth-inhibition mechanisms is triggering R-loop accumulation-induced genome instability and activating the cell-cycle checkpoint [27]. R-loops are relevant to neurodegenerative diseases and cancers, and they are also required for normal physiological process. Under normal conditions, R-loops are very rare in cells [28,29]. However, excessive R-loop accumulation is harmful to cells. C1orf109L-eGFP-interaction proteins are also involved in R-loop regulation. The immunofluorescence results showed that C1orf109L-eGFP shares co-location with the R-loops and immunoprecipitated R-loops with S9.6, an R-loop specific antibody; we then identified how C1orf109L-eGFP and its location on the R-loop can be significantly reduced by RNAse A treatment. We also found that C1orf109L-eGFP expression can trigger R-loop accumulation. Cell-growth inhibition caused by C1orf109L-eGFP expression can be partially reduced by overexpressing RNAseH1, an R-loop-specific digestion enzyme.

Although R-loop-related proteins were identified in HeLa cells, it is important to note that since cancer cells with abnormal epigenetic modification can cause many genes to exhibit low or no expression, it may lead to incomplete R-loop-related protein identification. Our work has now identified C1orf109L-eGFP as an RNA-processing regulator, which binds on the R-loop specifically and shares the same location with DHX9 (data not shown). In some abnormal cells, the epigenetic modification of *C1orf109*, such as in the promoter of *C1orf109* with hypermethylation, can cause lower *C1orf109* expression. We have also shown that *C1orf109* is not expressed in many kinds of cell lines, including HeLa cells. In HeLa cells, C1orf109 expression may not be related to the modification of DNA methylation and we did not detect acetylated histones of *C1orf109*; however, we did find that *C1orf109* mRNA and protein levels were increased with TSA treatment. It is possible that the lower expression of *C1orf109* may be related to histone acetylation modification in HeLa cells.

R-loops can cause cell-cycle arrest, thus inhibiting cell growth. We found that the cell-cycle arrested in the G2/M phase upon C1orf109L-eGFP expression. Furthermore, synchronizing the cell cycle at the G1/S boundary, and inducing C1orf109L-eGFP expression in different cell cycle phases, when C1orf109L-eGFPexpression in G1 phase, has shown that the cell cycle is impeded from the G1 to S phases. When C1orf109L-eGFP expression is in the S phase, the cell cycle has been impeded from the G2/M to the G1 phase. Interestingly, the S phase of HeLa cells only occurs within 7 hours; this means that C1orf109L-eGFP expression within a short time will impede the cell cycle from the G2/M phase to the G1 phase. Indeed, cell-cycle arrest occurs in the G2 phase. The R-loop regulates gene expression, such as the cell-cycle-related gene; however, expressing C1orf109 within a short time may not provide enough time to change the expression of other genes to alter the cell cycle. Therefore, arresting the cell cycle may not be related to the idea that C1orf109L-eGFP regulates cell-cycle-related gene expression.

Global gene-expression profiling revealed that there were no enriched cell-cycle functions. We also detected that there were no changes for G2 to M phase regulators, CDK1 and CCNB1 mRNA and protein levels, after releasing cell cycles 8 hours from G1/S boundary. As C1orf109L-eGFP caused cell-cycle arrest, it was primarily focused on R-loop-induced DNA damage. A chromosome spread assay suggested that C1orf109L-eGFP will cause a genome break; we also found unfolded chromosomes that belong to the G2 phase. The comet assay indicated that the expression of C1orf109 will cause DNA damage; further, the DNA double-stand break marker, γH2AX, had increased. Meanwhile, DNA damage that occurred downstream along the pathways was activated, which is reflected at the protein level of P21; moreover, inactivation phosphorylation of CDK1 had increased. When P21 was knocked down, cell growth reverted. These results indicate that C1orf109L-eGFP triggered R-loop accumulation and induced DNA damage to activate the cell-cycle checkpoint to inhibit cell growth; this means that gene expression regulation was not the main mechanism underlying this process.

Excessive R-loops can cause DNA damage; as such, promoting R-loop accumulation may be a novel way to induce cancer cell death in cancer therapy. CPT is a kind of anticancer medicine and an R-loop promoter that inhibits TOP1 to form DNA supercoils that cause the R-loop formation [30]. The aforementioned results indicate that C1orf109L-eGFP expression can bind with many RNA-processing proteins, such as DHX9, to promote R-loop formation. When CPT is used as a treatment, a loss of DHX9 promotes R-loop accumulation and DNA damage [31]. However, when SFPQ is knocked down to induce R-loop formation first, DHX9 will promote R-loop accumulation [32]. We found that C1orf109L-eGFP can trigger the R-loop to further increase in response to CPT to induce serious DNA damage. Meanwhile, the expression of C1orf109L-eGFP led to cell death in response to 5 hours of CPT treatment; further, the cell-death pathway was activated with CPT treatment in a time-dependent manner. However, we found that CPT treatment caused reduced fluorescence in the remaining surviving cells. It may be that the cell activated a protective mechanism to avoid C1orf109L-eGFP-induced DNA damage to promote cell death in response to CPT. We found that C1orf109L-eGFP began to degrade when CPT was used as a treatment for 4 hours, and fluorescence dramatically decreased at 8 hours. When treated with MG132, the degradation of C1orf109L-eGFP was significantly reduced. C1orf109L-eGFP can be degraded through the ubiquitinated proteasome pathway. Therefore, we found C1orf109L-eGFP ploy-ubiquitinated chain accumulation with CPT treatment, which demonstrated that CPT treatment promoted C1orf109L-eGFP degradation. This may be the cell’s protective system to avoid excessive R-loop formation to induced severe DNA damage. Meanwhile, we found C1orf109L-eGFP degradation when the cell cycle entered into the G0/G1 phase.

Therefore, we have partially uncovered the mechanism whereby C1orf109L-eGFP promotes cell-growth inhibition and genome instability caused by R-loop formation in human cells. We attempted to discover whether C1orf109L-eGFP can enhance R-loop-induced DNA damage and enhance chemosensitivity. We found that C1orf109L-eGFP can promote R-loop accumulation, and this induced severe DNA damage to activate cell death; however, there are other mechanisms that are involved in the induction of C1orf109L-eGFP degradation and which promote the cell cycle to enter into the G0/G1 phase, thus avoiding the cell death induced by CPT. In this study, we provided a promising target for tumour therapy and discussed feasibility and limitations for C1orf109L as target for tumor therapy. However, we did not exploit the inmost mechanism underlying how C1orf109L-eGFP triggers R-loop accumulation with or without CPT treatment. The inmost mechanism underlying this process, as well as that involved in C1orf109L-eGFP degradation in response to CPT, will be summarized in our next study.

## Methods

### Cell culture, antibody and reagents

HeLa, HEK-293 and HEK-293T cell lines were obtained from ATCC. The cells were grown in DMEM containing 10% fetal bovine serum (Biological Industries, BI) and 1% penicillin–streptomycin solution (Gibco) in a humidified incubator at 37℃ with an atmosphere of 5% CO_2_.

For antibody, mouse anti-Clorf109 monoclonal antibody was purchased from Genscript company (Nanjing, China.WB 1:1000 dilution), rabbit anti-eGFP polyclonal antibody(Abcam, WB 1:1000 dilution and IP:2 μg/ml, ab290), rabbit anti-PARP1 monoclonal antibody(Cell Signaling technology, WB 1:1000, Lot#9532), rabbit anti-phospho-Histone H2A.X(Ser139)(20E3) monclonal antibody(Cell Signaling technology, WB 1:1000 dilution, Lot#9718), rabbit anti-CDK1 polyclonal antibody(Proteintech, WB 1:2000 dilution, Catalog No:199532-1-AP), rabbit anti-CyclinB1 polyclonal antibody (Proteintech, WB 1:1000 dilution, Catalog No:55004-1-AP), rabbit anti-CyclinA2 polyclonal antibody(Proteintech, WB 1:1000 dilution, Catalog No:18202-1-AP), mouse anti-DHX9 polyclonal antibody (Abcam, WB 1:1000 dilution, ab26271), mouse anti-phospho-CDK1-T14 monclonal antibody(ABclonal, WB 1:1000 dilution, Catalog No:AP0015), rabbit anti-RNASEH1 polyclonal antibody(ABclonal, WB 1:1000 dilution, Catalog No:A9116), anti-GAPDH(ABclonal, WB 1:50000 dilution, AC033), mouse anti-S9.6 monclonal antibody (EMD millipore, IF 1:50 dilution, Lot#301193), mouse Flag-Tag monclonal antibody(Thermo Fisher, WB 1:1000 dilution, Catalog No: MA1-91878), rabbit anti-Histone H3ac(pan-acetyl) polyclonal antibody(Active Motif, WB 0.2μg/ml, Catalog No: 61637,61638), rabbit anti-Histone H3 polyclonal antibody(Proteintech, WB 1:1000, Catalog No:17168-1-AP), rabbit anti-NPM1(N-term) polyclonal antibody(Absin, WB 1:1000, abs106184)

For reagents, Trichostatin A(TSA)(MCE, Catalog No:HY-15144), Campathecin(MCE, Catalog No:HY-16560), MG132(Abcam, ab141003), Nocodazle(MCE, Catalog No:HY-13520), Doxycycline(MCE, Catalog No:HY-N05658), Phosphatase inhibitor cocktail 1(MCE, Catalog No:HY-K0022), protease inhibitor cocktail 1(MCE, Catalog No:HY-K0010) protein A/G magnetic beads (MCE, HY-K0202). Flag-M2 magnetic beads (theromo fisher, Lot:TJ276517)

### Plasmid constructs and siRNA

pTRIPZ-C1orf109L-eGFP, pLVSIN-RNaseH1-eGFP and pCMV-Flag-C1orf109L were constructed. Briefly, pLVSIN plasmid and pCMV-Flag were digested with EcoR I/Xho I at 37°C for 1 hour, the same condition were applied to digest pTRIPZ with Age I/Cla I, then desired digests were isolated by 1% agarose gel electrophoresis. The bands were excised and purified with Gel Extraction Kit (Theromo Fisher). For inserts, we get inserts from cDNA of HEK-293 by PCR. According to the need to construct the vector, inserts digest with the corresponding restriction endonuclease at 37°C for 1 hour, the desired inserts were purified with PCR Purification Kit (Theromo Fisher). The ligation reactions of insert with 50ng vector at a 3:1 molar ratio were performed at room temperature for 1 hour using T4-DNA ligase and buffer in a final volume of 10 μL. Ligation mixtures were used to transform the DH5α following the manufacturer’s instructions. 50 μL of transformation mixture were plated onto LB-agar containing ampicillin. 12 hours later, the selected clones and amplified with LB containing ampicillin. Finally, extract the plasmid from bacteria solution.

Ablation of p21 was performed by transfection with siRNA duplex oligos, which were synthesized by GenePharma Company (Shanghai, China). The sequences of the siRNAs and related primer were as supplementary table 1. Cell transfection was performed with Lipofectamine TM 2000 (Invitrogen) as described in the manufacturer’s protocol.

### Lentiviral Production

Human C1orf109L-eGFP Lentiviral Vector was constructed (pTRIPZ-C1orf109L-eGFP) and was used to overexpress human C1orf109L in HeLa and HEK-293. Viral particles were produced with a HEK-293T packaging cell lines, cells were infected once with viral supernatants. At day 2, infected cells were selected with puromycin for 7 days and placed in experiments. Overexpression of C1orf109L-eGFP was checked by Western blot and fluorescence inverted microscope (Olympus IX71).

### Protein extraction and western blotting

Proteins were extracted from sub-confluent cultures of cells and then characterized by western blot analysis. Cells were lysed in RAPI with phosphatase inhibitor cocktail, protease inhibitor cocktail, resolved on a sodium dodecyl sulphate-polyacrylamide electrophoresis(SDS) gel, and transferred onto an PVDF membrane (Millipore, Billerica, MA, USA). The membrane was blocked with 5% non-fat milk in phosphate buffer saline(PBS) containing 0.05% Tween-20 (PBST) for 1 hour at room temperature, and then probed with a primary antibody overnight at 4°C. After extensive washing, the membrane was incubated with a secondary antibody conjugated to horseradish peroxidase (1:10,000; Proteintech.) for 1 hour at room temperature. Blots were developed using ECL (Thermo Fisher Scientific, USA).

### Silver staining

After finished SDS-PAGE, we used Protein Silver Stain Kit stained the gel, Briefly, The procedure consisted soffixing with methanol, acetic acid and paraformaldehyde solutions, washed with ethanol (50% and 30%) and ddH_2_O, sensitizing with Na_2_S_2_O_3_.5H_2_O, washedwith ddH_2_O, impregnating with silver nitrate and paraformaldehyde solution, washed with ddH_2_O, developing with Na_2_CO_3_, paraformalde-hyde and Na_2_S_2_O_3_.5H_2_O solution, washed with ddH_2_O, and ending reaction with a stop solution-methanol 50%, and acetic acid 12%. Images were acquired with a camera (Cannon)

### RT-qPCR experiments

Total RNA was purified. Briefly, nucleic acids were extracted using Trizol reagent, Then, 1μg of total RNA was subjected to reverse transcription with random primers using Takera’s TranscriptorFirst Strand cDNA Synthesis Kit and cDNA was subjected to real-time PCR analyses using LightCycler 480 SYBR Green I Master Mix (Roche) in a LightCycler 480 Instrument(BD, Franklin Lakes, NJ). Relative expression levels quantifications were calculated by ΔΔCt method and normalized respect to GAPDH. Results were expressed as fold change over untreated condition.

### Tumour growth assay

Lentivirus-transduced C1orf109L-eGFP HeLa cells were suspended in 100 μL PBS and implanted subcutaneously into the flanks of nude mice (three 4- to 6-week-old male BALB/c nu/nu mice in each group; Laboratory Animal Unit, Harbin Medical University, China). Two weeks after inoculation, after tumor formation, nude mice with the same tumor size were divided into two groups: group I, DOX+(containing 2 mg/ml DOX 10 mg/ml sucrose water feed mice); group II, DOX-(containing 10 mg/ml sucrose water feed mice), tumors were calculated as follows: total tumor volume (mm^3^)=L×W^2^/2, where L is the length and W is the width. The sizes of the resulting tumors were measured per 2 days, the mice were sacrificed, and their tumors were dissected and weighed. Tissues were microscopically examined.

### Cell proliferation assays

HeLa-Tet-on and HEK-293-Tet-on cells were seeded into 96-well plates with a density of 5000 cells per well overnight (time 0) and treated with DOX or DMSO. After indicated time treatment, a mixed solution consisting of CCK-8(10 µL, MCE) and fresh culture medium (100 µL) was added to each well and incubated for an additional 2 hours at 37°C and 5% CO_2_. Finally, the absorbance at 450 nm was measured by a microplate reader (BioTek SynergyTM2). For colony formation assays, cells were seeded in six-well plate sat a density of 1000 cells per well treated with DOX or DMSO and cultured at 37°C for two weeks. After incubation, the cells were fixed with 100% methanol and stained with 0.1% (w/v) Crystal Violet. Capture pictures with camera (Cannon), the zoom picture acquire with stereoscopic microscope (Olympus SZX10)

### Flow cytometry analysis of cell cycle

Cells in different groups were trypsinized, washed once with PBS and fixed with 70% ethanol overnight at 4°C. After fixation, cells were washed once with PBS. After washing, cells were stained with PI/RNAse staining solution for 30 minutes (Tianjin Sungene Biotech. China). Flow cytometry (FCM) analysis was performed with a flow cytometer (BD Biosciences, USA).

For cell cycle synchronization, overnight post-plating, 2 mM Thymidine (TDR) was added to the culture medium. Following 16 hours incubation, cells were washed and fresh medium was added. Following 10 hours incubation, 2 mM Thymidine was again added to the culture medium and the cells incubated for a further 16 hours. Cells were then washed and contain DOX or DMSO media added. Following release from Thymidine block, for every indicated time, collect cell to detect cell cycle. And collect release time to 8 hours, protein and RNA of cells were extracted to detect the indicated genes expression.

### Immunofluorescence

HeLa-Tet-on were seeded with a density of 1×10^5^ per well in 12-well plates. After incubation for 24 hours, the cells were treated with DOX or DMSO and further incubated for 24 hours at 37°C and 5% CO_2_. Then, the cells on the slides were fixed with 4% PFA at room temperature for 10 minutes and permeabilized with 0.2% Triton X-100 at 37°C for 10 minutes, followed by incubation with anti-S9.6 antibody (dilution at 1:50) at room temperature for 60 minutes. Finally, the preparations were washed with PBS and mounted in fluorescent mounting medium with DAPI (Invitrogen). Negative controls were processed in the same way but without the primary antibody. Slides were photographed under a laser scanning confocal microscopy (Zeiss LSM510).

### Live cell imaging

Cells were plated on 35cm glass bottom dishes (MatTek). Imaging experiments were performed from 0 to 8 hours, for every 6 minutes collect a picture, with or without DOX induction for 24 hours. For CPT treatment experiment, HeLa-Tet-on C1orf109L-eGFP and HeLa-Tet-on RFP cells were plated on 35 cm glass bottom dishes with same number overnight. Treated cells with DOX and perform live cell imaging. 488nm(identify C1orf109L-eGFP expression cells) and 563 nm (identify RFP expression cells) channel were used. Imaging experiments were performed from 0 to 8 hours, every 6 minutes picture was collected. Image acquisition was performed using a live cell Imaging System (Observer Z1).

### Creating and analyzing protein-protein interaction networks

protein-protein interactions (PPI) among identified C1orf109L binding partners were extracted using the Search Tool for the Retrieval of Interacting Genes/Proteins (RRID:SCR_005223).The protein-protein interactions (PPI) was constructed by STRING website, only PPIs (*p*<0.001) from curated databases or curated published experiments were included in the PPI retrieval, and a minimum integrated confidence score of 0.5 was required for each interaction. Identified interactions were visualized using Cytoscape. We used the R-package “clusterprofiler” to perform GO annotation enrichment analysis, the *p* value were justified by the “Benjamini & Hochberg, BH” method [33]. The PPI network edge thickness represents the integrated confidence score for the interaction (ranging for medium confidence score of 0.5 to high confidence score of 1). Node colour in gray scale from light to dark indicates increasing abundance of the interactor in the C1of109L-eGFP immunoprecipitation, based on coverage rate of unique peptides (unique peptides number / length of protein) in LC-MS/MS data (ranging from 0.0007 to 0.006). Network and node level statistics were extracted from the resulting network using Cytoscape.

### Chromosome spreading

HeLa-Tet-on cells were treated with 200 ng/ml nocodazole for 16 hours. Cells were collected and swollen with 75 mM KCl for 20 minutes at 37°C. Swollen mitotic cells were collected and fixed with methanol: acetic acid (3:1). The fixing step was repeated two times. Cells were then dropped onto pre-hydrated glass slides and air-dried overnight. The following day, slides were mounted with Vecta shield medium containing DAPI. Images were acquired with a laser scanning confocal microscopy (Zeiss LSM510)

### Comet assay

After treatment with or without DOX for 24 hours, cells were collected for analyzing DNA damage activity by comet assay. Cell density was adjusted to 1000 cells mixed with 0.5% low-melting point agarose equilibrated to 37°C, and cell-agarose suspensions were spread onto the comet slides embedded with 1% normal-melting agarose for incubation at 4 °C. The prepared slides were lysed immediately in chilled neutral lysis buffer (0.1M Na_2_EDTA·2H_2_O, 2.5 M NaCl, 1% Triton X-100, 10% DMSO, and 10mM Tris, pH 8.0) at 4°C in the dark for 4 hours. In order to unwind the cellular DNA, the slides were submerged in the pre-cooled neutral electrophoresis buffer (90 mM Tris buffer, 90 mM boric acid, 2 mM Na_2_EDTA·2H_2_O, pH 8.5) at 4 °C for 20 minutes. Next, the damaged DNA fragments were electrophoresed for 30 minutes at a constant voltage of 20V·cm^−1^. After the slides being neutralized with 0.4 M Tris-HCl (pH 7.5) and left to air dry, samples were stained with the non-toxic DNA dye SYBR Green I at room temperature in the dark for 20 minutes and then examined under a fluorescence inverted microscope (Olympus IX71).

### Immunoprecipitation and DNA-RNA hybrid Immunoprecipitation(DR-IP)

For assay C1orf109L interaction protein, HeLa cells were cultured on 10cm dishes and grown to confluence. When cells at 60-70% confluence, transfect pCMV-Flag-C1orf109L for Flag tagged C1orf109L express for 36 hours. Cells were washed once with phosphate-buffered saline (PBS) and harvested by scraping and centrifugation at 800 g for 5 minutes. The harvested cells were washed with PBS and lysed for 30 minutes on ice in the lysis buffer (25mM Tris-HCl PH 8.0 250mM NaCl 1% Triton X-100) with Benzonase. Cell lysates were then spun down at 12,000 g for 20 min. The soluble fraction was collected, and the protein concentration was determined by Bradford assay. Next, 1 mg of extracted protein in lysis buffer was immunoprecipitated overnight with Flag-M2 affinity gel (Sigma) at 4°C. The immunoprecipitates were washed three times with lysis buffer.

The beads were then eluted with 0.5 mg/ml of the corresponding antigenic peptide for 4 hours or directly boiled in SDS loading buffer. For assay C1orf109L ubiquitination, breifly cells were lysed with lysis buffer (25 mM Tris-HCl pH 8.0, 250 mM NaCl, 1% Triton X-100, 1%SDS), and 100°C for 5 minutes then ultrasonication with 30% 5 seconds stop 5 seconds for 5min. 12,000 g for 20 min remove supernatant into a new tube and ten times dilution. The protein concentration was determined by Bradford assay. Next, 1 mg of extracted protein was immunoprecipitated overnight with GFP anti-body and mix with protein A/G magnetic beads at 4°C. The immunoprecipitates were washed three times with lysis buffer and boiled in SDS loading buffer.

For DR-IP, briefly, cells were isolated nuclear, suspend cells with nucleic extract buffer(10 mM Hepes pH 7.9, 1.5 mM MgCl_2_, 0.34 M sucrose, 10% glycerol, 1 mM DTT, 10 mM KCl) with 1% protease inhibitor, Triton X-100 (1%) was added and the suspensions were incubated for 5 min on ice. Nuclei were collected in pellet by 1300 g for 5 minutes at 4°C, then lysed nuclei with lysis buffer (25 mM Tris-HCl pH 8.0, 250 mM NaCl, 1% Triton X-100) with or without 1μg/ml RNAse A for 30 minutes and ultrasonication until DNA fragmant almost 200 bp. Finally 12,000 g for 20 minutes for supernatant, Supernatant was added S9.6 antibody for 4 hours. The immunoprecipitations were washed three times with lysis buffer and directly boiled in SDS loading buffer.

### Transcriptome analysis

Total RNA was extracted from sorted cells using an RNeasy MicroKit(Qiagen) following the manufacturer’s instructions. The RNA concentration and quality were assessed during the Qubit RNA assay kit (Invitrogen). We used 300 ng of total RNA to prepare the TruSeqlibrary, for which we used the Illumina Low-Throughput TruSeq RNA Sample Preparation Kit protocol, resulting inbarcoded cDNA. Next, 50 ng of barcoded TruSeq products were used for Illu-minaRNA sequencing on an IlluminaHiSeq2000 sequencer to generate single-end 50 or 51-nucleotide reads according to the manufacturer’s protocol. The expression levels of each sample were normalized as Reads Per Kilobase Per Million (RPKM) by dividing the read count of each transcript model with its length and scaling the total per sample to one million.

### Mass spectrometry analysis

Protein was added to a final concentration of 10 mM dithiothreitol (DTT), followed by final concentration 55 mM ammonium iodoacetate (IAM), finally added 1μg of Trypsin enzyme, overnight enzymatic hydrolysis 8 hours to 16 hours. The enzymatically produced polypeptide was desalted by a C18 column, and the dehydrated polypeptide was dried and dissolved in 15 μL of Loading Buffer (0.1% formic acid, 3% acetonitrile). The peptide was analyzed by LC-MS/MS (ekspertTMnanoLC; AB Sciex TripleTOF 5600-plus) instrument and the results were evaluated.

### Statistical analysis

All data were expressed in this manuscript as mean ± S.D. All the results have been performed at least three times by independent experiments. No samples and animals were excluded from the analysis. A two-tailed Student’s t test was used to analyze the statistical significance between two groups. The statistical analysis was performed by using GraphPad prism 7.0 (GraphPad Software Inc.). Asterisks indicate significant differences (\**p* < 0.05, \*\**p* < 0.01, \*\*\**p* < 0.001).

## Data availability

The data that support the findings of this study are available from the corresponding author upon reasonable request.

## Acknowledgements

This work was supported by the National Natural Science Foundation of China (No.31571323 and No.305400669.)

## Author contributions

Y.L conceived this study; D.P and L.Yq designed the study; D.P and X Wq. performed the experiments; D.P and Z. Xq performed the animal experiments; Z. Xh and L. Yq performed the mass spectrometry analyses; H.F and D.P performed the data statistics. D.P wrote the manuscript with comments from all authors.

